# A global dataset of seaweed net primary productivity

**DOI:** 10.1101/2021.07.12.452112

**Authors:** Albert Pessarrodona, Karen Filbee-Dexter, Kira A. Krumhansl, Pippa J. Moore, Thomas Wernberg

**Affiliations:** UWA Oceans Institute and School of Biological Sciences, University of Western Australia, Crawley, Western Australia 6009, Australia; Institute of Marine Research, His, Norway; Fisheries and Oceans Canada, Bedford Institute of Oceanography, Dartmouth, Nova Scotia Canada; School of Natural and Environmental Sciences, Newcastle University, Newcastle-Upon-Tyne, NE1 7RU, UK; Department of Science and Environment, Roskilde University, Universitetsvej 1, DK-4000 Roskilde, Denmark

## Abstract

Net primary productivity (NPP) plays a pivotal role in the global carbon balance, but estimating the NPP of underwater habitats remains a challenging task. Seaweeds (marine macroalgae) form the largest and most productive underwater vegetated habitat on Earth. Yet, little is known about the distribution of their NPP at large spatial scales, despite more than 70 years of local-scale studies being scattered throughout the literature. We present a global dataset containing NPP records for 242 seaweed species at 419 individual sites distributed on all continents from the intertidal to 55 m depth. All records are standardized to annual aerial carbon production (g C m^-2^ yr^-1^) and are accompanied by detailed taxonomical and methodological information. The dataset presented here provides a basis for local, regional and global comparative studies of the NPP of underwater vegetation, and is pivotal for achieving a better understanding of the role seaweeds play in the global coastal carbon cycle.

## Background and Summary

NPP is a major driver of ecological functioning and a key flux in the global carbon cycle ^1^. The advent of remote sensing technologies has facilitated the measurement of terrestrial ^2–4^, freshwater ^5,6^, and oceanic ^7,8^ NPP at unprecedented scales, with most global models of NPP available to date relying on space-based observations ^4,6^. In contrast, the magnitude, patterns and determinants of spatial and temporal variation of primary productivity in the coastal ocean remains poorly understood ^9^. This is particularly true for submerged vegetated habitats, where most observations rely on *in situ* measurements due to remote-sensing measurements being challenged at shallow depths ^10,11^. Existing measurements of coastal vegetation NPP vary in methodology and are usually reported in different units, hindering our understanding of the role these habitats play in the carbon cycle and how it compares to other primary producers^12^. Additionally, the majority of measurements are conducted at local scales, which means compilation of multiple local-scale datasets is required to unravel larger spatiotemporal patterns ^11^.

Seaweeds form the largest and most productive underwater vegetated habitat on Earth, drawing a flux of CO2 comparable to the Amazon rainforest every year (Duarte et al. 2021). The carbon assimilated through this production fuels local marine food webs ^13,14^ and can constitute a trophic subsidy to areas with low primary production such as soft-bottom communities ^15^. Recent studies also suggest that seaweed carbon makes important contributions to oceanic carbon export ^16^, with some estimates highlighting seaweeds as major contributors to oceanic carbon sequestration ^17^. This has reopened the debate on their potential use as carbon dioxide removal and/or climate change mitigation tools ^18,19^, although great uncertainties exist in the carbon fluxes they underpin ^17^. Indeed, despite the fact that it has been more than 70 years since seaweeds were shown to be amongst Earth’s most productive organisms ^20–22^, we still know little about how their NPP varies across taxa, space and time ^23^. Previous attempts to collate seaweed NPP data at large spatial scales have been geographically restricted (e.g.^12,24^) or focused on specific taxa (e.g. ^25,26^). These limitations have precluded a global understanding of the patterns and determinants of NPP across seaweed taxa, which is in urgent need to inform on the promising potential of seaweeds.

Here we describe the most comprehensive global dataset of marine macroalgae NPP gathered to date. Data was obtained from the primary literature or provided directly by authors and contains records from a total of 242 taxa from 419 sites in 72 different ecoregions. Measurements of seaweed NPP were collected at the taxa level, and reflect per-area productivity rates across a range of depths and seaweed groups. Each record is accompanied by detailed descriptions of methodology used and is classified into habitat groups depending on the growing substrate, vegetation height and dominant vegetation at the study site.

## Methods

### Data compilation

An extensive search of published reports, PhD thesis and the peer-reviewed literature was performed to capture studies dealing with the net primary productivity or biomass production of marine macroalgae. First, a formal search was performed in the Scopus database using the search terms “primary AND product* OR growth or npp AND (seaweed OR alga* OR kelp OR rocky AND reef OR turf OR temperate AND reef OR coral OR polar OR Arctic)”, which yielded 470 entries (September 2020). We then filtered the query by searching for relevant content in the title and abstract, yielding a total of 60 studies. Further searches were conducted in the China National Knowledge Infrastructure database (CNKI), J-STAGE repository (Japan) and Scientific Electronic Library Online (SciELO) to capture studies (with English abstracts) from underrepresented regions such as Asia and South America. Additional studies were included from existing reviews on the productivity of tropical ^27,28^, temperate ^12,29^ and polar algae ^30^ and from being cited in the scanned papers. Finally, we included a few more studies from MSc or PhD thesis, the authors’ unpublished data, and other published reports based on our knowledge of the research field.

### Data selection and quality control

Given that our analysis was centered on patterns of annual areal carbon production by seaweeds, each of the potentially relevant studies was then evaluated against the following set of criteria to determine if they could be included in the final dataset. First, studies had to examine seaweed NPP or biomass accumulation on a per area basis. This criterion excluded studies examining biomass-specific productivity rates (e.g. ^24,31^) unless those rates were applied to standing biomasses or covers in the field (e.g. ^32^). Second, studies had to provide estimates of NPP at the primary producer level with minimal interference of heterotrophic organisms. This criterion excluded studies examining net ecosystem primary production (NEP) and metabolism, which usually rely on diel dissolved oxygen measurements in the water column (mostly applied in coral reefs, e.g. ^33,34^) or directly above the benthos (e.g. Aquatic Eddy Covariance method ^35,36^). Third, studies had to capture seasonal variability in NPP across the year. This criterion excluded studies conducted at a single point in time, month or season, with the exception of studies concerning annual species where the growth or biomass accumulation was measured at the end of their life-cycle (i.e. the maximum period of growth). Fourth, quantification of productivity had to be performed *in situ* on the reef or outdoor mesocosms mimicking natural reef conditions. This criterion excluded laboratory-only experiments, aquaculture yields, models (e.g. Ecopath models) and field studies in which the natural environmental conditions were experimentally modified (e.g. nutrient enrichment, acidification, sediment additions). Fifth, details of the specific sampling location and measuring method had to be provided. Sixth, studies had to provide new data not previously reported in other publications. After applying the criteria above, our final filtered dataset featured 1,048 records from 229 independent studies published between 1967 and 2020 and covering a range of seaweed vegetation types (Fig. 1a, b).

**Figure 1.**
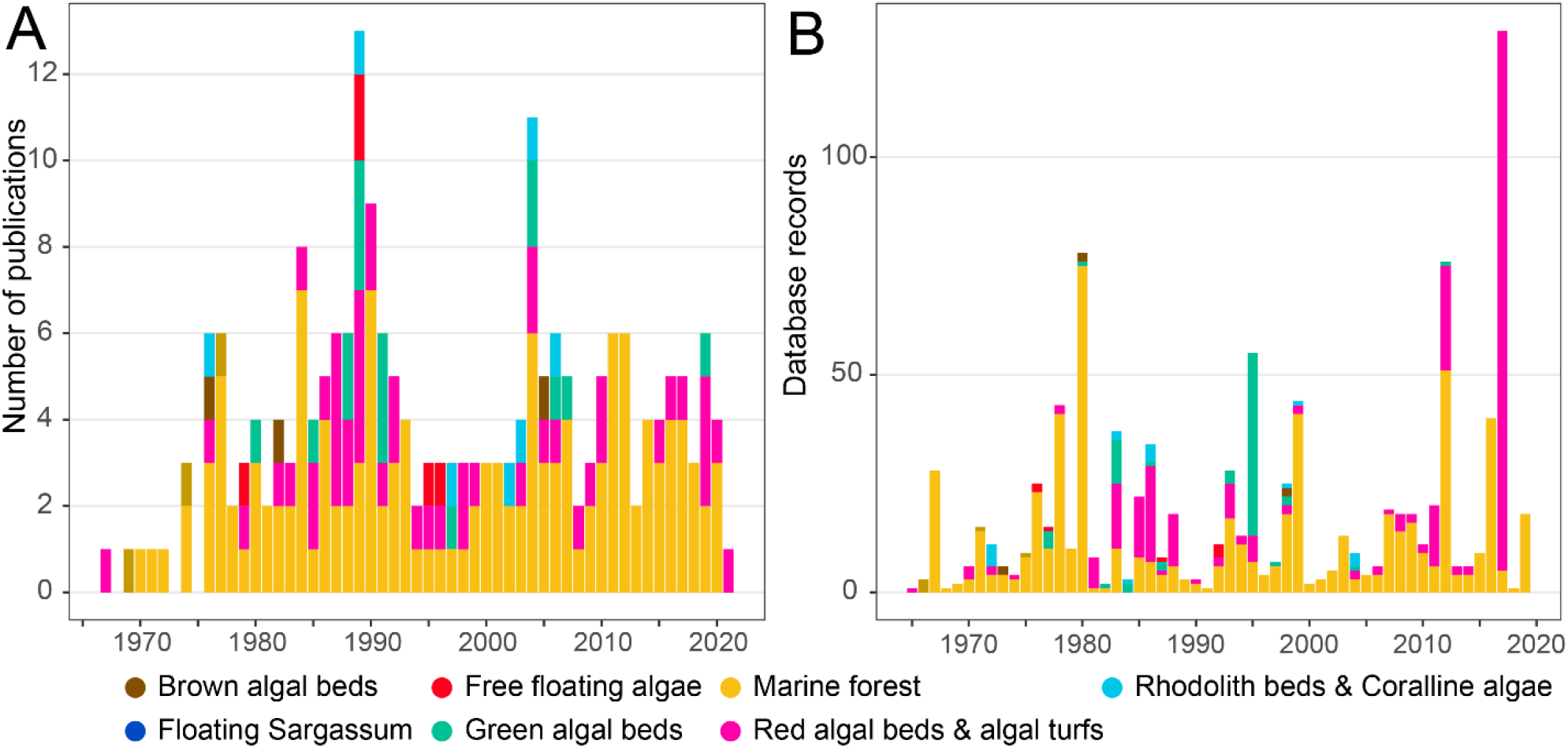
Temporal coverage of seaweed NPP measurements conducted at different habitat types and by tidal level (intertidal or subtidal), which are indicated in different colours. (a) Number of database records (i.e. a measurement of NPP per taxa, depth, site, year and method) depending on when the measurements were conducted (i.e. not published). (b) Number of studies by date of publication.

Available data were extracted into an excel template from the suitable articles’ text, tables, figures (using the graph digitizing tool *Webplot Digitizer* ^37^) or supplementary material. In our study, a record was considered to be the aerial net primary productivity of a taxon over the course of a year. If the data in a given study was not directly reported as annual rates, these where computed based on the monthly, bimonthly or seasonal means, with the corresponding standard deviation also being computed. The sampling effort (frequency of measurements throughout the year) was also recorded as it may have impacted the estimates’ accuracy. Data were entered into the template in the same units as the original source, but were also standardized to annual areal carbon production (i.e. g C m^-2^ y^-1^). Values reported in fresh or dry weight (FW, DW respectively) were converted to carbon using species- and genera-(most cases) or family- and order-specific factors when these were not available for a given species. Conversion factors provided in the studies were preferably used, but otherwise these were derived from the database provided in ^38^. Metadata describing the depth, substrate, sampling year and season, taxonomy, study site and its geolocation, measuring method and data extraction procedure was attached to each individual row. When a given value was not available, this is entered as “NA”. If a study reported NPP from multiple taxa, depths, sites, methods or time points, these were entered as separate case studies (separate rows). NPP of taxa within the same sample plot (e.g. multi-species *Sargassum* bed, kelp and understory algae) was also entered as separate records, but a specific column was created to denote that data would require summation of the rows to yield total areal productivity.

A site was defined as a single location where NPP was measured using the criteria above, with its geographic coordinates being added as metadata. If these were not directly provided in the article, we used the maps and/or description of the study locations to approximate their coordinates on Google Maps, noting also that these were approximations in the record’s metadata. NPP records across depths were considered to be within the same study site as long as measurements were within 30 m of each other. Each independent site was given a unique ID within each study.

As different sampling methods measure different aspects of photosynthesis and carbon assimilation^12^, we also recorded the method used to estimate each value of NPP. These were grouped into several subcategories which fell into two basic approaches: photorespirometry and biomass accumulation (Table 1). Photorespirometry-based methods measure direct carbon assimilation, while biomass accumulation measures only the carbon destined to plant growth, and thus is expected to always yield lower estimates of NPP ^39^. An overview and discussion of the advantages disadvantages of each method is provided elsewhere (e.g.^12,40^).

**Table 1.** Summary of the methods to estimate seaweed NPP in our database.

| General Method | Method | Description | Examples |
| --- | --- | --- | --- |
| Biomass accumulation | Single Harvest | Production is assumed to be equal to the maximum standing biomass after a period of time. Production can be estimated by outplanting tiles into the field and quantifying their algal biomass in a given timeframe, or by harvesting annual species when they reach the end of their life cycle | 43,44 |
|  | Periodic Harvest | Periodic harvests of standing biomass over short time scales. Changes in standing biomass are attributed to growth or losses. Production can be estimated by subtracting the maximum and minimum biomass achieved, summing of all positive increments, or by counting individuals of a cohort and their mean weight through time (Allen method) | 45–47 |
|  | Commercial harvest | Periodic harvests of standing biomass but targeting certain vegetative structures. Plants are not cultured but rather grown on the reef | 48,49 |
|  | Tagging | Individual-plant increases in weight are followed through time by tagging, staining or punching holes in the plant. The mean individual increases in biomass are then multiplied by plant density to obtain areal rates | 50,51 |
| **Photo-respirometry** | Gas evolution (in situ) | Measurements of dissolved oxygen (or more rarely CO<sub>2</sub>) of individuals or communities enclosed in transparent benthic changes. Measures true NPP (carbon assimilation) by subtracting gross primary productivity to respiration. Respiration rates obtained by enclosing individuals in dark chambers | 52,53 |
|  | Gas evolution (mesocosm) | Measurements of dissolved oxygen in individuals maintained in outdoor mesocosms with flow through seawater and field-like levels of irradiance | 54,55 |
|  | Gas evolution (modelling) | Relationship between photosynthesis and irradiance established, and photosynthesis modelled based on irradiance changes throughout the year | 56 |
|  | Isotopes | Thalli are submerged in water enriched with isotopes and uptake by macroalgal tissue is measured after a given period of time. Measures true NPP as well as carbon isotope tracers (<sup>14</sup>C or more rarely <sup>13</sup>C) | 57,58 |

Studies and taxa were also classified according to the habitat where measurements were performed using the information given within the published article (Table 2). Habitat categories were defined based on key structural parameters like vegetation height, the dominant vegetation (e.g. brown, red or green algae) as well as their position within the water column (benthic or pelagic). Within a study, taxa from different groups could be classed in the same habitat (e.g. canopy, epiphytes and understory algae all being part of a “marine forest”) unless they formed distinct patches within the habitat matrix (e.g. red algal bed patches interspersed with marine forests ^41,42^), or the study examined different depth bands, sites or habitats. When incubations of different taxa were performed in isolation within a study, these were independently assigned a habitat category.

**Table 2.** Definitions for the habitat type category. Categories were based on vegetation height, dominant vegetation (brown, red or green algae) as well as their position in the water column (benthic or pelagic).

| Habitat type | Description | Examples |
| --- | --- | --- |
| Marine forest | Vegetation dominated by large canopies formed by brown algae from the orders Laminariales, Fucales, Tilopteridales and Desmarestiales. Includes understory and epiphytic taxa associated with the canopies. | Kelp & *Sargassum* forests |
| Brown algal beds | Low-lying vegetation dominated by brown algae | *Padina* beds |
| Red algal beds & algal turfs | Low-lying vegetation dominated by red algae and/or algal turfs, which contain aggregations of multiple species. | *Gelidium*, *Gracilaria* beds, algal turfs |
| Green algal beds | vegetation dominated by green algae, including *Halimeda* biohermes, green-tides both in subtidal and intertidal areas | *Caulerpa* beds, *Halimeda* bioherme |
| Rhodolith beds & coralline algae | Habitats coralline algae forming rhodolith beds, from tropical and temperate areas | Coralline barrens |
| Floating Sargassum | Pelagic *Sargassum* rafts (*S. fluitans*, *S. natans*) | *Sargassum* rafts |
| Other floating algae | Other free-floating aggregations of algae on the bottom or the sea surface | Ulva blooms |

## Data records

The dataset is publicly accessible for download in the Figshare repository (https://doi.org/10.6084/m9.figshare.14882322.v1).

### Taxonomic coverage

The database contains NPP information for 242 species or taxonomic entities (e.g. crustose coralline algae, algal turf), from 49 families, 26 orders and all major seaweed groups and functional forms. The majority of species with NPP records are brown algae (55 %; kingdom Phaeophyta) (Fig. 2a), with just over half the database being composed of records from the orders Laminariales and Fucales (558 records, Fig. 2b)

**Fig 2.**
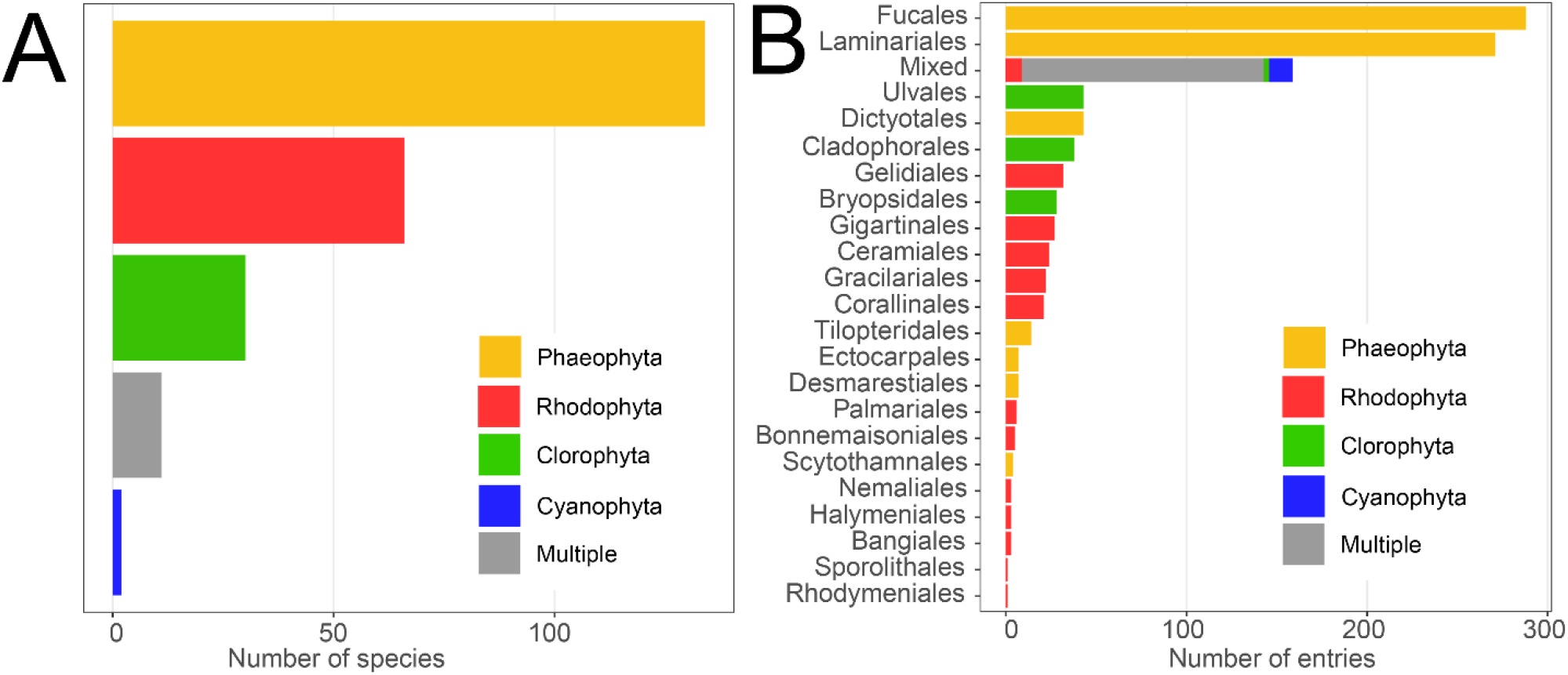
Taxonomic coverage of the database. Multiple denotes taxonomic groupings that involve species from different phyla (e.g. algal turfs).

### Spatial and temporal coverage

The dataset contains NPP data from 419 sampling sites (Fig. 3a) spanning from the high intertidal level (3 m above mean sea level mark) down to 55m (Fig. 3b). Sites span all major oceanographic realms and are distributed form the poles to the tropics, with the majority of records concentrated in temperate latitudes 40-60° and concerning marine forests. The vast majority of studies measured NPP over 1-2 years. Only 2% of records report measurements conducted ≥ 3 years, and only three records report continuous NPP measurements >10 years. The temporal resolution of the measurements conducted within the sampling period varies from biweekly to annual measurements.

**Fig. 3.**
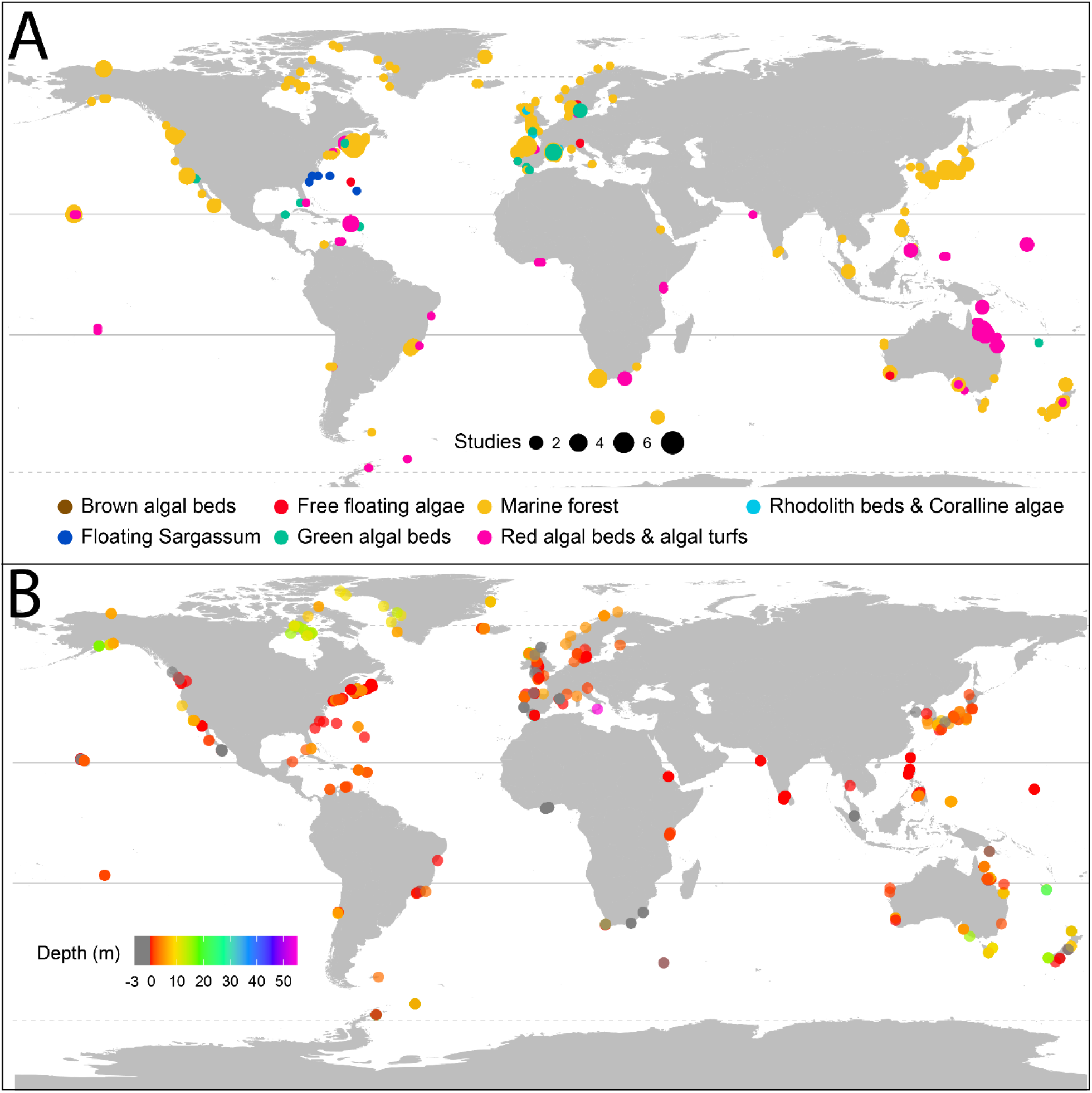
Location and depth of the 421 study sites included in the database. (a) Each circle represents were. Study depth of the records in our database. Measurements conducted in the intertidal (i.e. above sea level are indicated in grey).

### Data collection sources and methods

Records were mostly extracted from the literature (94%), followed by unpublished personal data (2.5%), PhD and MSc thesis (2.5%) and a minor fraction corresponding to published reports in the grey literature. Most of the data was sourced from tables and text (73% of records), whilst the rest was extracted from graphs (20%) or from raw data. The vast majority of NPP records in the database were obtained using biomass-accumulation-based methods (87%), followed by photorespirometry-based methods (12.9%), with only a tiny fraction of records using both methods. While these two methods measure different aspects of carbon assimilation, NPP patterns from both methods are largely consistent across latitude (Fig. 4). Biomass accumulation measurements are well distributed globally (Fig. 5a), while photorespirometry-based measurements are common in coral reefs (mostly on algal turfs), pelagic Sargassum spp. rafts and a few other temperate locations (Fig. 5b).

**Figure 4.**
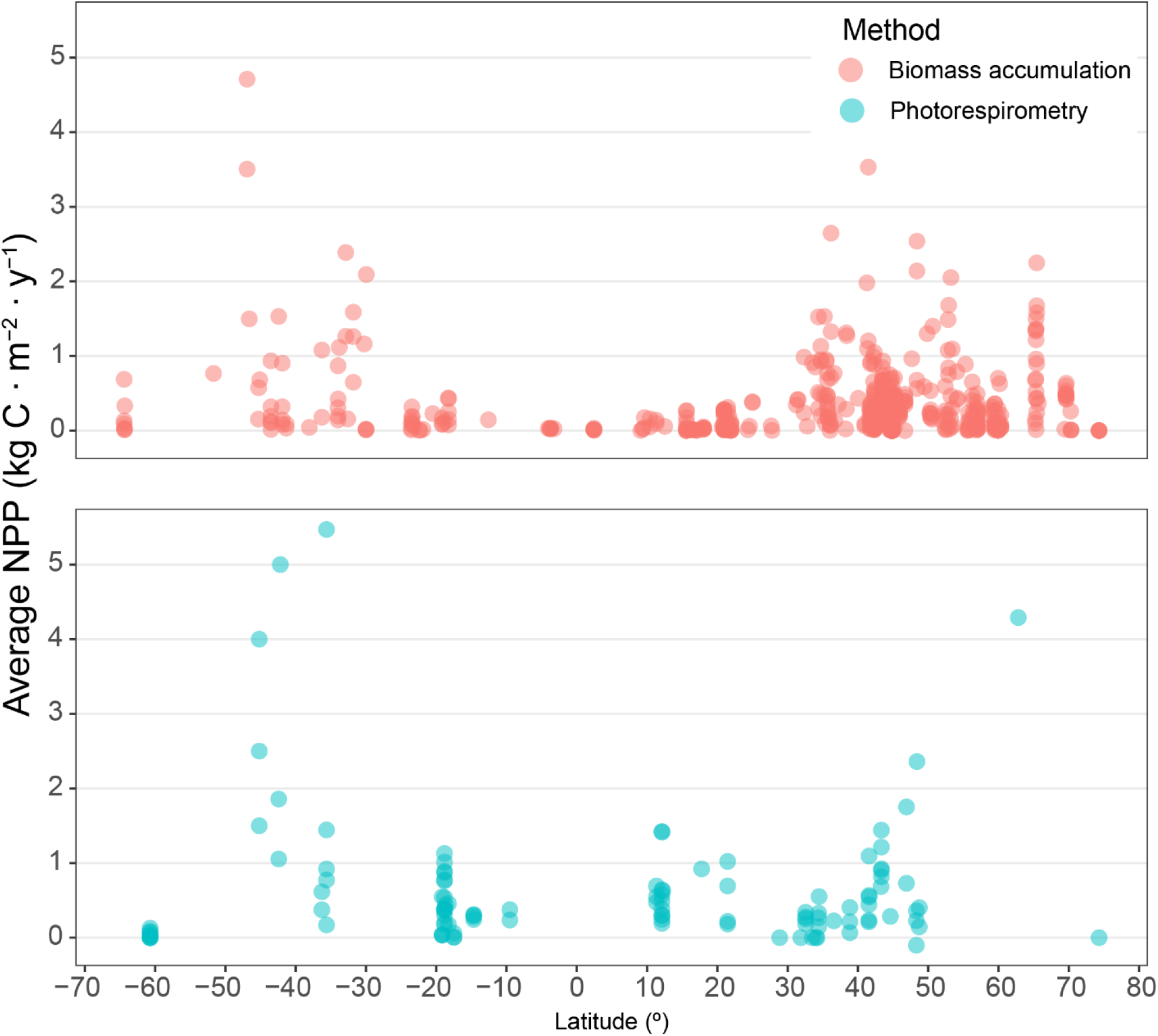
Latitudinal patterns of observed NPP depending on measuring methods. Dots indicate the average NPP of a study conducted within a given location.

**Fig. 5.**
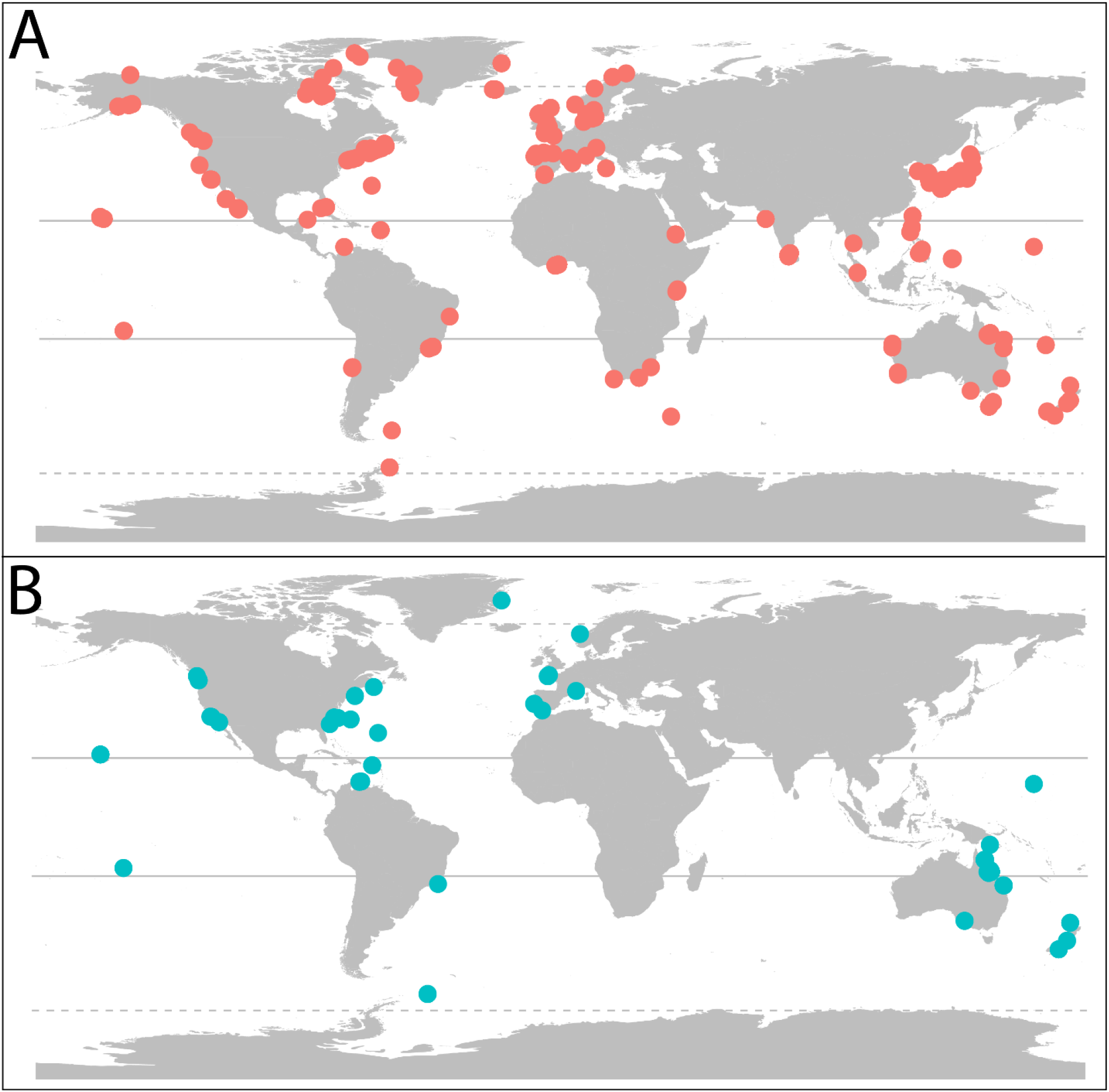
Distribution of observations depending on the broad methodology to measure NPP. A) Biomass-based accumulation and B) bhotorespirometry-based methods.

## Technical Validation

The database was curated by the authors, and each of the records entered is associated with their initial in the “Person_entering_data” column. We used templates to minimize the entry of spelling errors, inconsistencies, and incorrect values. Upon finalizing data entry, we conducted quality control by

i. Checking taxonomic names. The validity of taxa names was checked using the taxon match tool of the World Register of Marine Species (WoRMS) in May 2021. The names were corrected and updated if taxonomies had changed since publication of the study.
ii. Checking geographic coordinates. We projected the coordinates on a 1:10,000,000 shapefile of the world’s landmasses (EPSG:3857) checking they did not lay on land.
iii. Checking for duplicates. Records with identical NPP values for the same species and GPS coordinates were double checked for accuracy.
iv. Checking for outliers. Frequency histograms and quantile plots were generated to evaluate potential outliers. Records with very small (<1 gC m^-2^ y^-1^, i.e. 10% quartile) or large (<1,100 gC m^-2^ y^-1^, i.e. 95% quartile) NPP values were double checked for accuracy.

## Usage Notes

Each of the records (rows) in our database provides the average annual aerial NPP and standard deviation (when reported) of a given taxon at a given site, depth, year and study and by a given measuring method. Each record is also accompanied by a series of metadata describing the taxonomic information, geographic coordinates, description of the measuring method used and vegetation and substrate type. The variables’ (columns) definitions and descriptions can be found in Table 3. When the taxa measured includes species from multiple genera, families, orders or classes, this is indicated as “Mixed”.

Despite our efforts to obtain measurements across the globe, our dataset contains taxonomical and geographical biases (Fig. 2,3), with most records concerning brown algae and few records being available from South America, Africa, the Indian Ocean and Antarctica. We advise that researchers using the database should be aware of the influence these biases might have on their analyses.

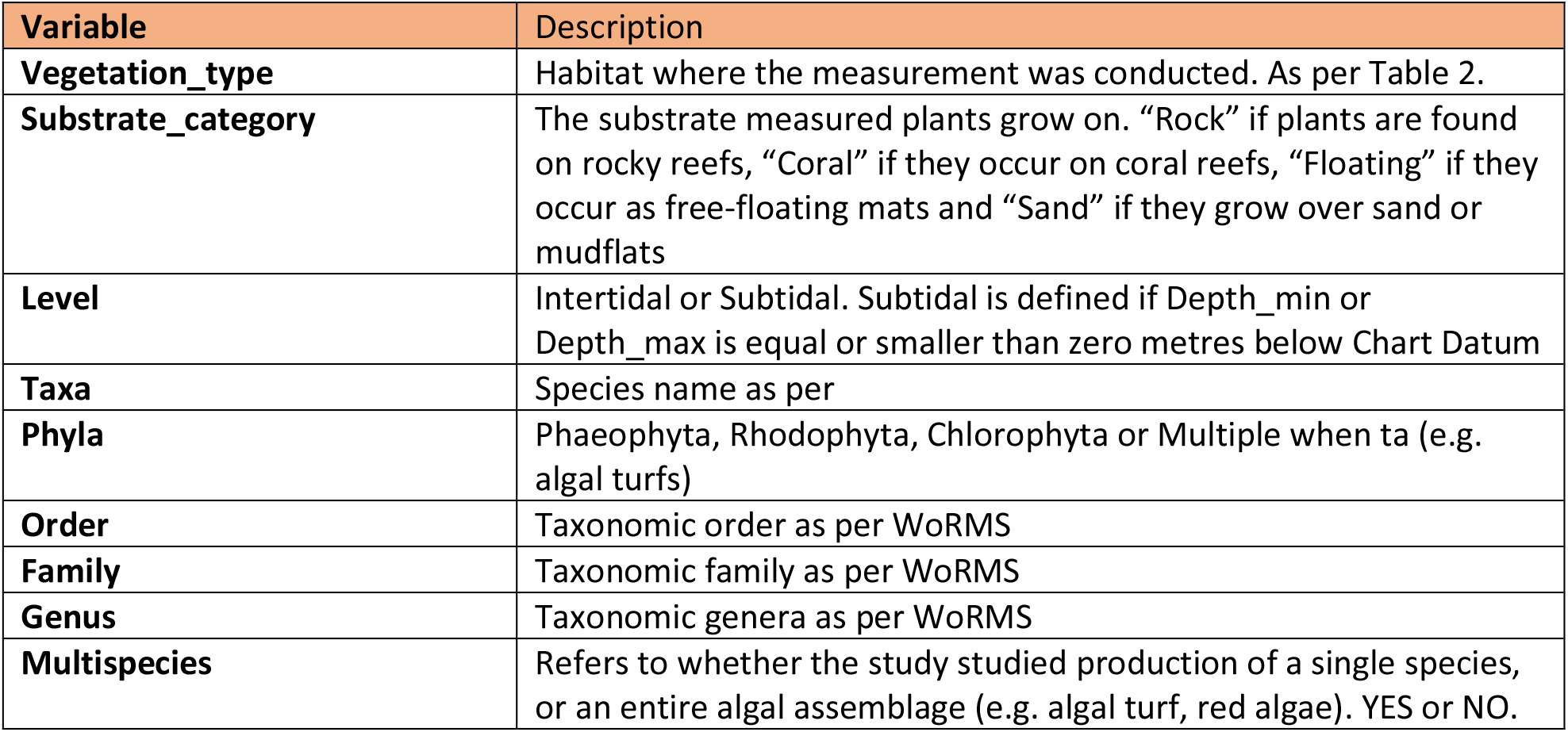
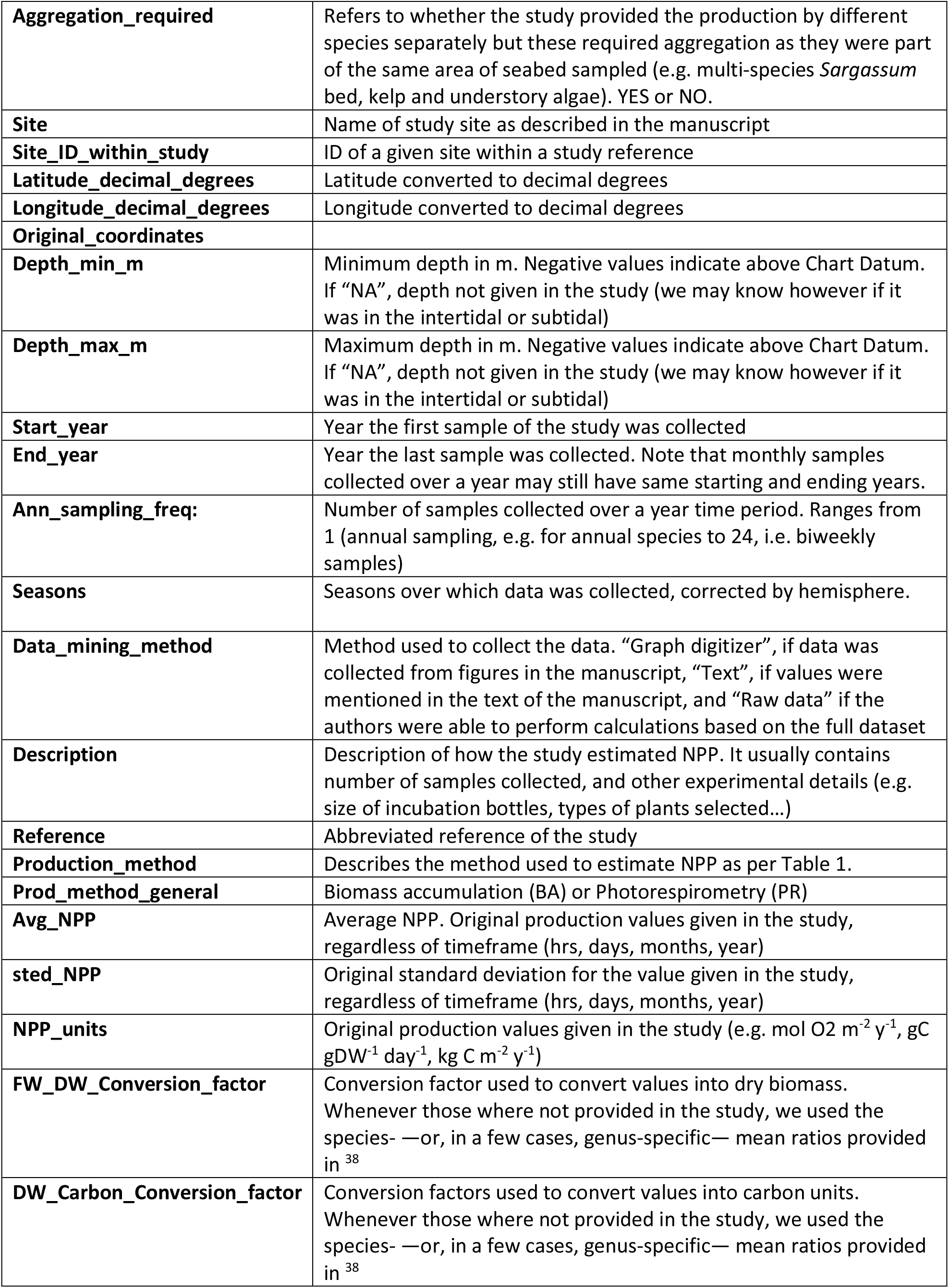
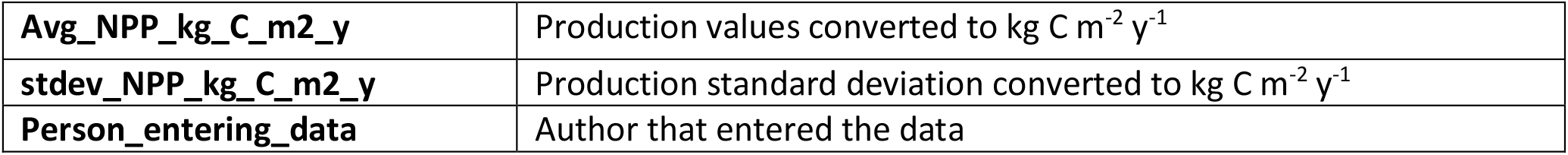

| Variable | Description |
| --- | --- |
| **Vegetation_type** | Habitat where the measurement was conducted. As per Table 2. |
| **Substrate_category** | The substrate measured plants grow on. “Rock” if plants are found on rocky reefs, “Coral” if they occur on coral reefs, “Floating” if they occur as free-floating mats and “Sand” if they grow over sand or mudflats |
| **Level** | Intertidal or Subtidal. Subtidal is defined if Depth_min or Depth_max is equal or smaller than zero metres below Chart Datum |
| **Taxa** | Species name as per |
| **Phyla** | Phaeophyta, Rhodophyta, Chlorophyta or Multiple when ta (e.g. algal turfs) |
| **Order** | Taxonomic order as per WoRMS |
| **Family** | Taxonomic family as per WoRMS |
| **Genus** | Taxonomic genera as per WoRMS |
| **Multispecies** | Refers to whether the study studied production of a single species, or an entire algal assemblage (e.g. algal turf, red algae). YES or NO. |
| **Aggregation_required** | Refers to whether the study provided the production by different species separately but these required aggregation as they were part of the same area of seabed sampled (e.g. multi-species *Sargassum* bed, kelp and understory algae). YES or NO. |
| **Site** | Name of study site as described in the manuscript |
| **Site_ID_within_study** | ID of a given site within a study reference |
| **Latitude_decimal_degrees** | Latitude converted to decimal degrees |
| **Longitude_decimal_degrees** | Longitude converted to decimal degrees |
| **Original_coordinates** |  |
| **Depth_min_m** | Minimum depth in m. Negative values indicate above Chart Datum. If “NA”, depth not given in the study (we may know however if it was in the intertidal or subtidal) |
| **Depth_max_m** | Maximum depth in m. Negative values indicate above Chart Datum. If “NA”, depth not given in the study (we may know however if it was in the intertidal or subtidal) |
| **Start_year** | Year the first sample of the study was collected |
| **End_year** | Year the last sample was collected. Note that monthly samples collected over a year may still have same starting and ending years. |
| **Ann_sampling_freq:** | Number of samples collected over a year time period. Ranges from 1 (annual sampling, e.g. for annual species to 24, i.e. biweekly samples) |
| **Seasons** | Seasons over which data was collected, corrected by hemisphere. |
| **Data_mining_method** | Method used to collect the data. “Graph digitizer”, if data was collected from figures in the manuscript, “Text”, if values were mentioned in the text of the manuscript, and “Raw data” if the authors were able to perform calculations based on the full dataset |
| **Description** | Description of how the study estimated NPP. It usually contains number of samples collected, and other experimental details (e.g. size of incubation bottles, types of plants selected...) |
| **Reference** | Abbreviated reference of the study |
| **Production_method** | Describes the method used to estimate NPP as per Table 1. |
| **Prod_method_general** | Biomass accumulation (BA) or Photorespirometry (PR) |
| **Avg_NPP** | Average NPP. Original production values given in the study, regardless of timeframe (hrs, days, months, year) |
| **std_NPP** | Original standard deviation for the value given in the study, regardless of timeframe (hrs, days, months, year) |
| **NPP_units** | Original production values given in the study (e.g. mol O<sub>2</sub> m<sup>-2</sup> y<sup>-1</sup>, gC gDW<sup>-1</sup> day<sup>-1</sup>, kg C m<sup>-2</sup> y<sup>-1</sup>) |
| **FW_DW_Conversion_factor** | Conversion factor used to convert values into dry biomass. Whenever those were not provided in the study, we used the species- —or, in a few cases, genus-specific— mean ratios provided in <sup>38</sup> |
| **DW_Carbon_Conversion_factor** | Conversion factors used to convert values into carbon units. Whenever those were not provided in the study, we used the species- —or, in a few cases, genus-specific— mean ratios provided in <sup>38</sup> |
| **Avg_NPP_kg_C_m2_y** | Production values converted to $\text{kg C m}^{-2} \text{ y}^{-1}$ |
| **stdev_NPP_kg_C_m2_y** | Production standard deviation converted to $\text{kg C m}^{-2} \text{ y}^{-1}$ |
| **Person_entering_data** | Author that entered the data |

